# Lateral recurrence dynamically refines visual representations for task performance

**DOI:** 10.64898/2026.10.02.756219

**Authors:** Amirhossein Farzmahdi, Hossein Adeli, Wang Boran, Chase King, Nikolaus Kriegeskorte

**Author notes:** Equal contribution (Amirhossein Farzmahdi, Hossein Adeli).

## Abstract

Recurrent connections are pervasive in visual cortex, and visual representations evolve as recurrent processing unfolds. Yet the mechanism underlying this temporal transformation remains poorly understood. To address this question, we trained recurrent convolutional networks on face and object recognition and tested them on a parametric stimulus set in which category, identity, and pose were systematically varied. This design isolates how lateral recurrence reshapes neural-population representational geometry and single-unit tuning across recurrent timesteps. Within the trained domain, recurrence increased the separability between identities in deeper layers, while leaving the non-trained domain representations largely unchanged. Gating lateral weights abolished the across-timestep gain in trained-domain identity decoding, demonstrating that lateral recurrence drives this refinement. At the single-unit level, recurrence strengthened the correlation between each unit’s category preference and its within-domain identity sensitivity, providing a unit-level mechanism for the population-level changes. These results identify lateral recurrence as a dynamic mechanism for refining learned representational structure within a fixed neural population, yielding testable predictions for time-resolved recordings in domain-selective cortex.

## Introduction

The primate ventral visual pathway transforms retinal input into representations that support object categorization and individual identification. This transformation has traditionally been modeled as a feedforward hierarchy of increasingly invariant feature detectors (Riesenhuber and Poggio, 1999; DiCarlo and Cox, 2007). Yet the ventral stream is not purely feedforward: lateral connections within areas and feedback projections between areas are anatomically pervasive and give rise to neural dynamics beyond the initial feedforward sweep (Felleman and Van Essen, 1991; Lamme and Roelfsema, 2000; Gilbert and Li, 2013). Recurrent interactions have been implicated in diverse computations including contextual modulation, predictive inference, and perceptual grouping (Rao and Ballard, 1999; Lee and Mumford, 2003; Roelfsema, 2006). Recurrent connectivity structure can enable a neural network to achieve greater and more flexible computational depth using limited hardware, to implement iterative algorithms, to integrate priors and priorities into visual inference through expectation and attention, and to exploit sequential dependencies in the data (van Bergen and Kriegeskorte, 2020). As a result of recurrent signal flow, the representations enabling recognition are not fixed after the feedforward pass but evolve over time (Xie et al., 2025; Kietzmann et al., 2019; Anthes et al., 2026).

Our working hypothesis here is that lateral connectivity dynamically *refines* the representations, shaped by visual experience and task demands. The nature of this refinement, however, remains unresolved: recurrence could leave the feedforward representation unchanged (Hypothesis A, static), refine representations equally across domains (Hypothesis B, domain-general), or refine them selectively for certain domains (Hypothesis C, domain-specific) (Fig. 1E). It is also possible that recurrent refinement of representations has both domain-general and domain-specific components.

**Figure 1.**
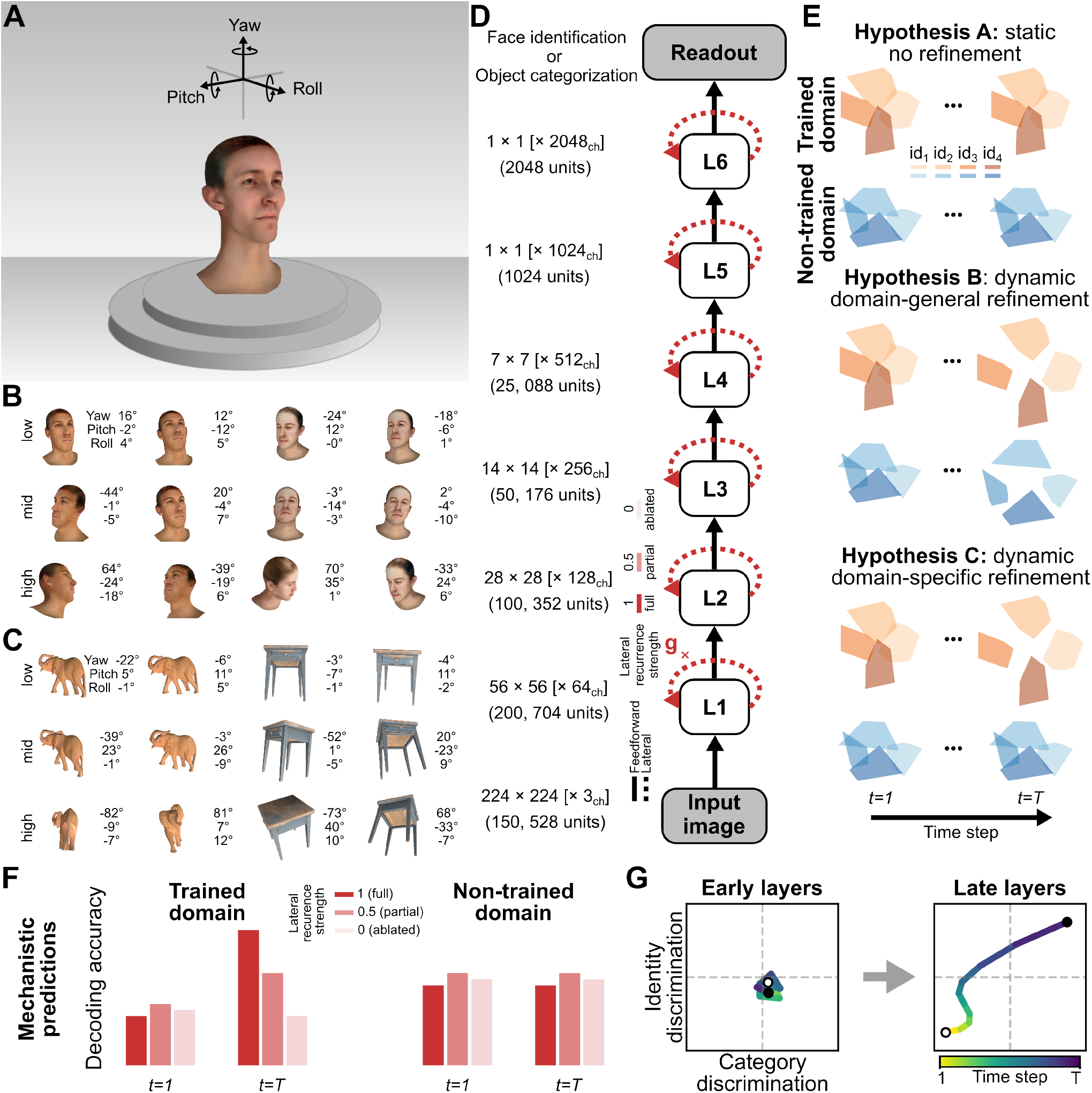
Stimulus design, model architecture, and competing hypotheses. (**A**) Parametric 3D stimulus generation for faces and objects. Each identity is rendered under controlled variations in yaw, pitch, and roll, with illumination and camera settings held constant within each domain. (**B**) Example face stimuli at three pose-variation levels (low, mid, high). Numbers indicate the corresponding yaw, pitch, and roll angles. (**C**) Example object stimuli generated using the same pose-variation scheme as the face stimuli. (**D**) Recurrent convolutional network architecture. The model consists of six hierarchical layers (L1–L6) with feedforward (solid) and lateral recurrent connections (dashed). Spatial resolution and channel dimensionality are indicated for each layer; a linear readout is applied to the final layer. The lateral recurrence strength can be gated by a scalar gain factor *g*, allowing causal manipulation of lateral recurrence (*g* = 1 intact recurrence; *g* = 0 ablated recurrence). (**E**) Three competing hypotheses for how lateral recurrence acts on task-relevant representations established by the feedforward sweep. Under static recurrence (Hypothesis A), the representational geometry produced by the feedforward sweep is preserved across recurrent timesteps in both the trained and non-trained domain, with no further refinement of identity structure. Under dynamic domain-general refinement (Hypothesis B), recurrence reshapes representational structure over time in both domains equally, regardless of training history. Under dynamic domain-specific refinement (Hypothesis C), recurrence expands between-identity distances in the trained domain across timesteps, while leaving non-trained-domain representations mainly unchanged. Orange and blue denote trained and non-trained domains, respectively; shaded polygons depict identity-level representations at the initial state (*t* = 1) and after recurrent processing (*t* = *T*). (**F**) Predictions under dynamic domain-specific refinement: decoding accuracy should increase across timesteps for the trained domain and scale with lateral recurrence strength, while the non-trained domain shows comparatively little benefit. (**G**) Predicted temporal evolution in a two-dimensional space defined by category discrimination (*x*-axis) and identity discrimination (*y*-axis). Lower layers are expected to show limited changes across recurrent timesteps, whereas higher layers should follow a stronger trajectory toward enhanced identity discrimination. Color indicates recurrent timesteps.

Under the domain-general refinement hypothesis (Hypothesis B), recurrence acts as a general-purpose mechanism that enhances representations, regardless of category. For example, recurrent dynamics in domain-general, lower-level stages might reduce noise or shrink the representation toward a prior for natural inputs. Recurrent circuits have been shown to strengthen feature representations under degraded input conditions such as occlusion, noise, and blur (Wyatte et al., 2012; Spoerer et al., 2017; Tang et al., 2018; Rajaei et al., 2019; Kar et al., 2019), and disrupting recurrent processing by backward masking can render objects invisible (Fahrenfort et al., 2007). Predictive coding frameworks propose that feedback signals sharpen the tuning of lower-level neurons largely independent of stimulus domain (Rao and Ballard, 1999; Lee and Mumford, 2003). Under this view, recurrence functions as a domain-general sharpening process, most useful when the feedforward estimate is uncertain.

The domain-specific refinement hypothesis (Hypothesis C) posits that recurrence refines representations in a manner that depends on the category of the image. For example, recurrence in higher-level category-specific circuits could use domain-specific prior knowledge to refine or regularize the representation. In the primate face-processing system, identity representations emerge alongside category selectivity and gain pose-tolerance across hierarchical stages (Freiwald and Tsao, 2010; Chang and Tsao, 2017). Task-optimized feedforward networks also develop domain-specialized representations resembling those observed in cortex (Dobs et al., 2022). Perceptual learning studies show that training selectively enhances discriminability for practiced distinctions, not all stimulus features (Schoups et al., 2001; Jehee et al., 2012; Goldstone, 1998). These findings establish that the feedforward sweep delivers task-relevant representations. How recurrence transforms the representations over time, whether by domain-general or domain-specific refinement, remains unclear in part because the development of the primate visual system is interactively determined by neuroanatomy, visual experience, and task training.

Task-optimized deep neural network models provide a useful framework for disentangling the contributions of architecture, visual diet, and task demands. Networks trained end-to-end on large-scale recognition tasks account for key aspects of ventral-stream representations (Khaligh-Razavi and Kriegeskorte, 2014; Yamins et al., 2014; Kubilius et al., 2018; Schrimpf et al., 2018; Doerig et al., 2023), and recurrent architectures better capture the temporal dynamics of human and primate visual processing (Kietzmann et al., 2019; Kar et al., 2019; Spoerer et al., 2020). Because architecture and recurrent connectivity structure can be held fixed while the training objective is varied, these models enable us to ask how learned recurrent interactions reshape representations through repeated layer updates. A recurrent network can be unrolled and written as a feedforward computation with shared weights. However, the updates from recurrent connections are applied to the same units, so the neural population stays fixed while its representation evolves over time. This framework is well-suited for tracking how population geometry and single-unit tuning change across recurrent timesteps.

Here, we ask specifically how *lateral* recurrence transforms representations over time. Previous studies suggest that lateral connections play an important role in within-area computations such as contextual modulation, surround interactions, and contour integration (Gilbert and Li, 2013; Angelucci et al., 2017). This makes lateral recurrence a natural candidate mechanism for reshaping representations within a fixed neural population over time.

We focus on representations of faces, whose representation at the category and identity level has been amply investigated with neural recordings in macaques (Tsao et al., 2006; Freiwald and Tsao, 2010) and functional imaging in humans (Puce et al., 1995; Kanwisher et al., 1997; Haxby et al., 2001; Gobbini and Haxby, 2006; Kriegeskorte et al., 2007; Anzellotti et al., 2014; Nestor et al., 2011; Schiltz and Rossion, 2006). We use object representations as a control domain to investigate to what extent dynamic transformations learned through task training are domain-specific.

We compare models with identical recurrent convolutional architectures, trained for either face or object recognition, and evaluate them using a controlled parametric stimulus set in which category, identity, and pose variation are systematically varied. This design separates coarse-scale category representations from fine-scale identity representations, enabling us to test how each changes across timesteps. We track these changes using cross-pose decoding to assess the tolerance of the identity representations to pose variation. We also characterize single-unit selectivity and neural-population representational geometry. Finally, we probe the role of lateral recurrent connections through graded ablation, using gating of lateral weights to assess their causal contribution to the dynamic computations.

Our results show that lateral recurrence builds on the task-specific organization present after the feedforward sweep and progressively reorganizes it. In the trained domain (and especially in deeper layers) recurrence preserves the coarse category structure established by the feedforward sweep, while expanding the representational volume devoted to distinguishing entities within the trained category with robustness to variation in pose. Non-trained-domain representations, by contrast, remain largely unchanged.

At the single-unit level, this refinement gradually enhances the alignment between category preference and within-domain identity sensitivity over recurrent timesteps: The units that are most strongly tuned to the trained category also become the units that best discriminate identities within it. The learned lateral weights selectively enhance trained-domain identity information over time.

## Results

We compared lateral-recurrent convolutional networks with identical architectures but different training objectives. Each model was trained on either face recognition or object recognition (Fig. 1D). We tracked how category and identity information evolved across layers and recurrent timesteps using linear decoding and representational geometry analyses (see Methods).

### Recurrence selectively enhances trained-domain identity decoding

We trained cross-validated linear decoders on the final layer (L6) to discriminate identities across changes in pose (see Methods). In the face-trained network, identity decoding for faces increased across recurrent timesteps, whereas decoding for object-identity remained flat (Fig. 2A). The object-trained network showed the opposite pattern: object-identity decoding improved over recurrent time, whereas face-identity decoding remained flat (and near chance; Fig. 2B). This double dissociation cannot be explained by any property of the stimuli, which were held fixed across the two models: the same object images that the object-trained model refines are left largely unchanged by the face-trained model, and the reverse holds for the face images. A domain-general refinement (Hypothesis B) would instead predict recurrent gains in both domains within each model. Instead, recurrence selectively transforms trained-domain representations over time: pose-tolerant decodability of identity information grew for the trained domain, while it remained unchanged for the non-trained domain. A model trained jointly on both objectives learned lateral connectivity that increased pose-tolerant decodability of identity for both domains across time, with recurrence refining identity information aligned with each unit’s preferred domain (Supplementary Fig. S1).

**Figure 2.**
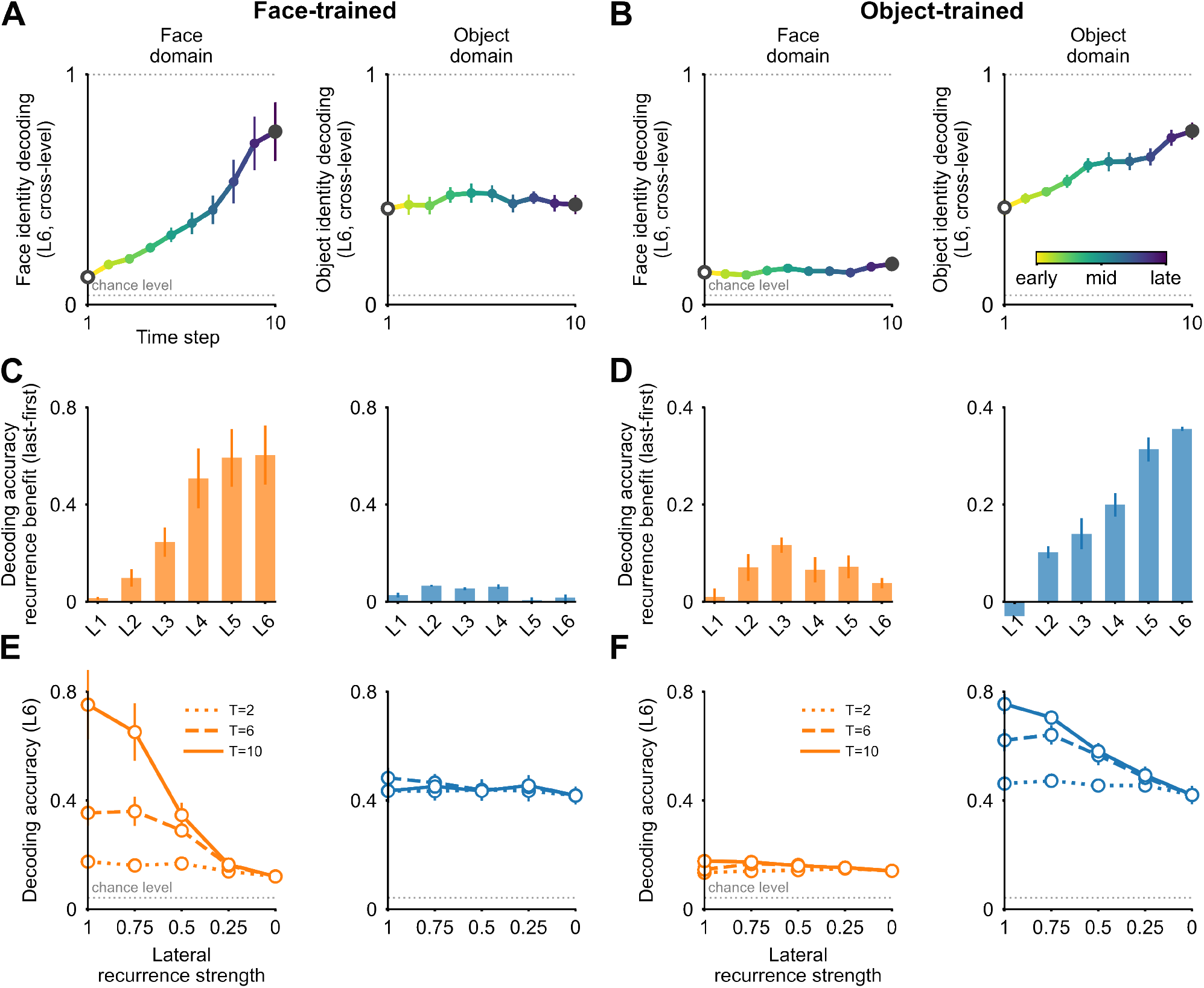
Recurrent processing selectively improves identity decoding in the trained domain. (**A**) Cross-pose identity-decoding accuracy at layer L6 across recurrent timesteps in the face-trained model, using leave-one-level-out generalization across the three pose-variation levels (low, mid, high). Color indicates recurrent timestep from early to late (light to dark). Face-identity decoding rises well above chance, whereas object-identity decoding remains nearly flat. Error bars indicate SEM across leave-one-level-out pose splits (see Methods). (**B**) Cross-pose identity-decoding accuracy at layer L6 across recurrent timesteps in the object-trained model. Object-identity decoding improves markedly across timesteps, whereas face-identity decoding remains near chance, yielding the complementary pattern seen in panel A. (**C**) Recurrence gain (decoding accuracy at the last timestep minus the first timestep) for the face-trained model across layers L1–L6, shown separately for the face domain (orange; left) and object domain (blue; right). Substantial gain emerges in mid-to-late layers (L4–L6), primarily in the face domain. Error bars indicate SEM across the three leave-one-level-out pose splits (n = 3). (**D**) Recurrence gain for the object-trained model across layers. The pattern reverses: mid-to-late layers show pronounced gain primarily in the object domain. (**E**) Parametric variation of lateral recurrence strength in the face-trained model at L6, from full strength (1) to full ablation (0), shown for three recurrent durations (*T* = 2, dotted; *T* = 6, dashed; *T* = 10, solid). Face-identity decoding (left) declines monotonically as recurrence strength decreases, with stronger dependence at longer recurrent durations, whereas object-identity decoding (right) remains largely unchanged. Error bars indicate SEM across leave-one-level-out pose splits. (**F**) Same parametric analysis for the object-trained model. Object-identity decoding (right) shows a graded dependence on recurrence strength that becomes stronger with longer recurrent durations, whereas face-identity decoding (left) remains largely unchanged.

This selectivity also depended on depth within the model hierarchy. The recurrence benefit, defined as the difference in decoding accuracy between the last and first timesteps, was negligible in lower layers but pronounced in mid-to-high layers (L4–L6), and confined to the trained domain (Fig. 2C–D). Temporal refinement was therefore concentrated in deeper layers. This does not imply that recurrence is unimportant in early visual areas. In the present task, however, recurrent changes were concentrated where higher-order feature combinations and identity-relevant structure are most likely to be represented, consistent with the role of inferotemporal cortex in complex object representations (Hu et al., 2025).

To test whether this selectivity depends directly on lateral recurrence, we performed a graded ablation experiment. We parametrically gated recurrence strength from full (1) to completely ablated (0). Reducing recurrence strength monotonically impaired decoding in the trained domain, with larger effects at later timesteps, whereas decoding in the non-trained domain was largely unaffected (Fig. 2E–F). The graded dose–response confirms that lateral recurrence is necessary for the accumulation of identity information across timesteps, especially in deeper layers. All decoding patterns reported in this section were reproducible across independent training runs with different random initializations (Supplementary Fig. S2).

We next asked whether these decoding gains reflect changes in representational geometry across recurrent time.

### Recurrence reshapes late-layer geometry toward identity refinement

The decoding results show that identity information becomes increasingly accessible over recurrent timesteps in the trained domain. However, decoding alone does not reveal how the underlying population representation is reorganized. We therefore examined the full geometry of population responses by computing representational dissimilarity matrices (RDMs) for each layer and recurrent timestep (Fig. 3).

**Figure 3.**
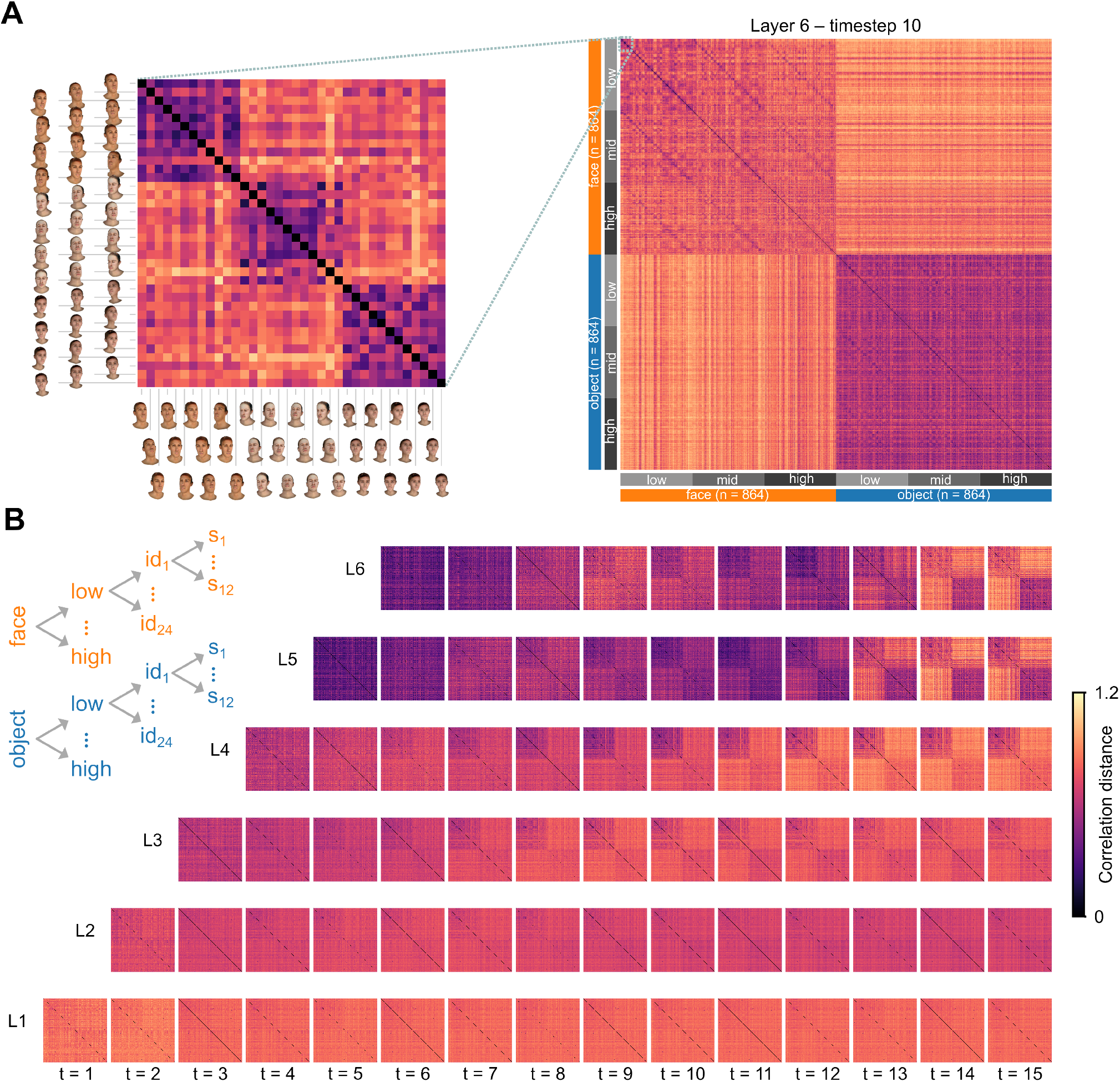
Population geometry evolves across layers and recurrent timesteps. All results shown are from the face-trained model. (**A**) Representative representational dissimilarity matrix (RDM) from the final layer (L6) at the last recurrent timestep (t = 10). The schematic (left) shows a 36 × 36 sub-block—3 identities × 12 samples within the low pose-variation level—illustrating how within-identity comparisons (small distances, near-diagonal blocks) are distinguished from between-identity comparisons (larger distances). The full RDM (right) shows pairwise correlation distances between all 1,728 stimulus representations, organized by domain (face, object), then pose-variation level (low, mid, high), then identity (24 identities per domain, 12 samples per identity); lighter colors indicate larger (more dissimilar) distances. A coarse block structure separates faces from objects (category), while finer structure along the diagonal and para-diagonals within each domain reflects identity discrimination. (**B**) RDMs across all layers (L1, bottom; L6, top) and recurrent timesteps, shown left to right from early to late. Under the staggered unrolling scheme, higher layers receive their first input later and therefore have fewer available timesteps (see Methods). In the highest layers, the face block sharpens its diagonal and para-diagonal (identity) block structure over recurrent time, while the object block remains comparatively undifferentiated.

In the higher layers of the face-trained model, recurrent processing increased the average dissimilarity between different individual faces while decreasing the average dissimilarity between different poses of the same face. Responses to different poses of the same identity became more similar, whereas responses to different identities became more separated. This reveals how recurrence improved linear decodability: it reorganized the geometry, making it more similar to a pose-tolerant code for identity. Object representations in the same model showed comparatively little reorganization. The object-trained model showed the opposite pattern: recurrence expanded the clusters corresponding to object identities. The geometric transformation, thus, tracked the trained objective rather than acting uniformly across domains.

To quantify these temporal changes across the full model hierarchy, we computed second-order distances between RDMs from every combination of layer and timestep (Fig. 4A). Multidimensional scaling of the second-order distance matrix reveals each layer’s representational dynamics as a trajectory through the space of representational geometries (Lange et al., 2022), as well as the relationships between layer representations. All representational geometries fall on a smooth 2D manifold within the space of RDMs whose dimensions correspond to layer (vertical) and time (horizontal, Fig. 4B). The 2D RDM manifold is well represented in the flat MDS map (Pearson correlation coefficient between 2D Euclidean distances and inter-RDM Spearman correlation distances: 0.97). Lower layers make smaller adjustments overall to their representational geometries over timesteps than higher layers (shorter trajectories). Lower layers also tend to make bigger adjustments early on and only small adjustments in the final timesteps: representational dynamics slow toward convergence. The final layer L6 underwent the most profound representational transformation.

**Figure 4.**
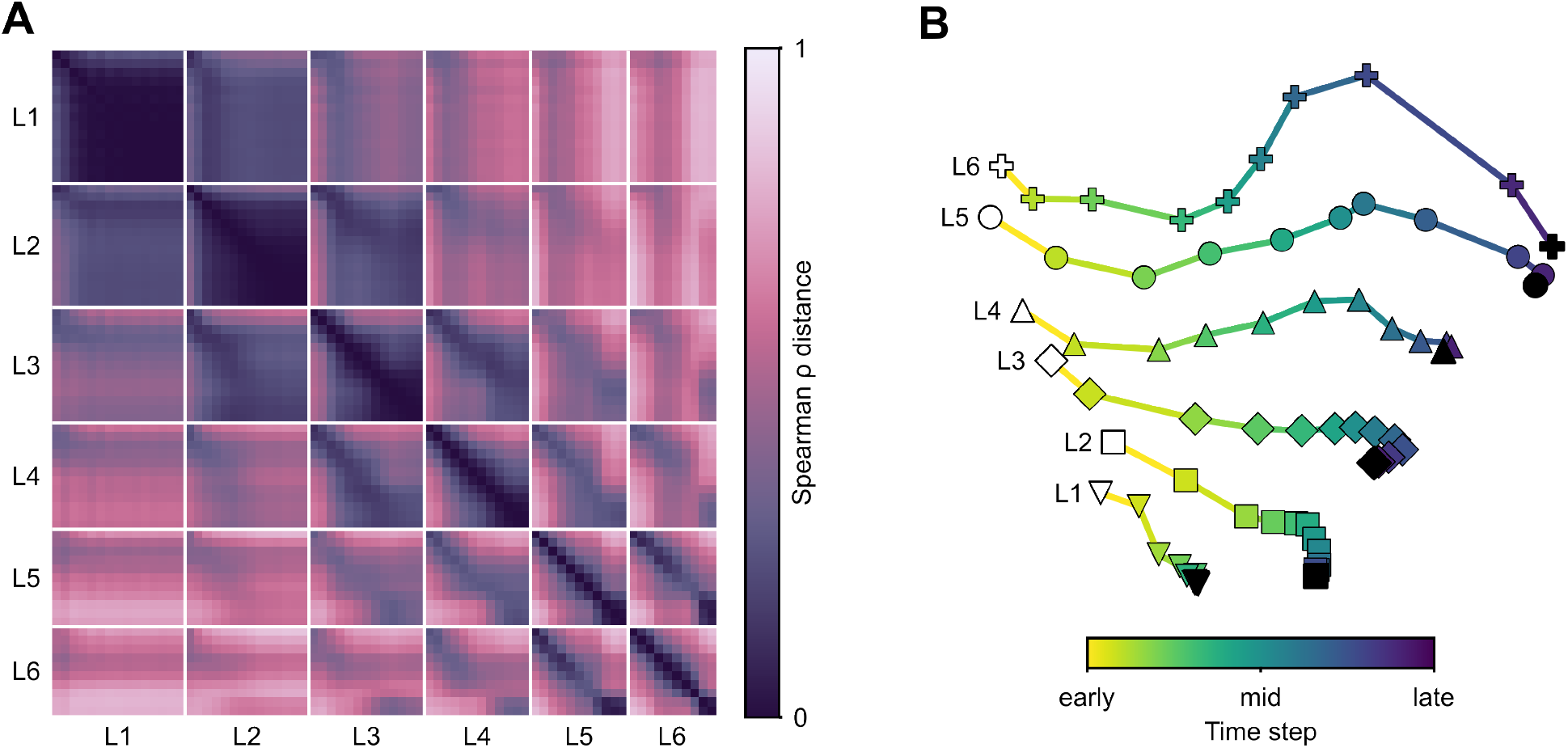
Representational change accumulates in higher layers across recurrent time. All results shown are from the face-trained model. (**A**) Second-order representational dissimilarity analysis. Each cell shows the Spearman *ρ* distance between a pair of RDMs derived from specific layer–timestep combinations. Within-layer progression across recurrent timesteps produces graded geometric shifts in higher layers, whereas lower layers remain relatively stable. Off-diagonal structure indicates that high-layer representations at later timesteps become increasingly dissimilar from both low-layer and early-timestep states. (**B**) Multidimensional scaling (MDS) embedding summarizing the second-order distances between layer–timestep representational states shown in panel A. Each point represents a unique layer–timestep state, with marker shape indicating layer and color indicating timestep (yellow, early; dark purple, late). Lower layers (L1–L3) form tight clusters with minimal temporal displacement, whereas mid-to-high layers (L4–L6) traverse substantially longer representational distances across recurrent time, with L6 showing the largest displacement. This pattern was replicated using normalized Euclidean distance (Supplementary Fig. S3).

These results show that recurrence yields a complex and highly structured pattern of transformations of the population geometry across layers and time. Representational dynamics were most profound in the higher layers, which also showed the strongest recurrence benefits for trained-domain decoding. We next tested which dimensions of this geometry were most affected by recurrence: coarse category structure and finer within-category identity structure.

### Category structure stabilizes early and pose-tolerant identity representations emerge later

We derived clustering indices from the RDMs to separately quantify category clustering (faces vs. objects) and within-category identity clustering (Fig. 5A). The category-clustering index measured how much larger the average between-category dissimilarity was than the average within-category dissimilarity (in units corresponding to the average within-category dissimilarity). The identity-clustering index measured how much larger the average between-identity dissimilarity was than the average within-identity dissimilarity (in units corresponding to the average within-identity dissimilarity). Note that a high clustering index implies linear discriminability, but not vice versa: Two classes can have perfect linear discriminability and arbitrarily low clustering if their response distributions are thin sheets that are close together like stacked pancakes. Clustering, thus, imposes a stronger requirement than mere linear discriminability.

**Figure 5.**
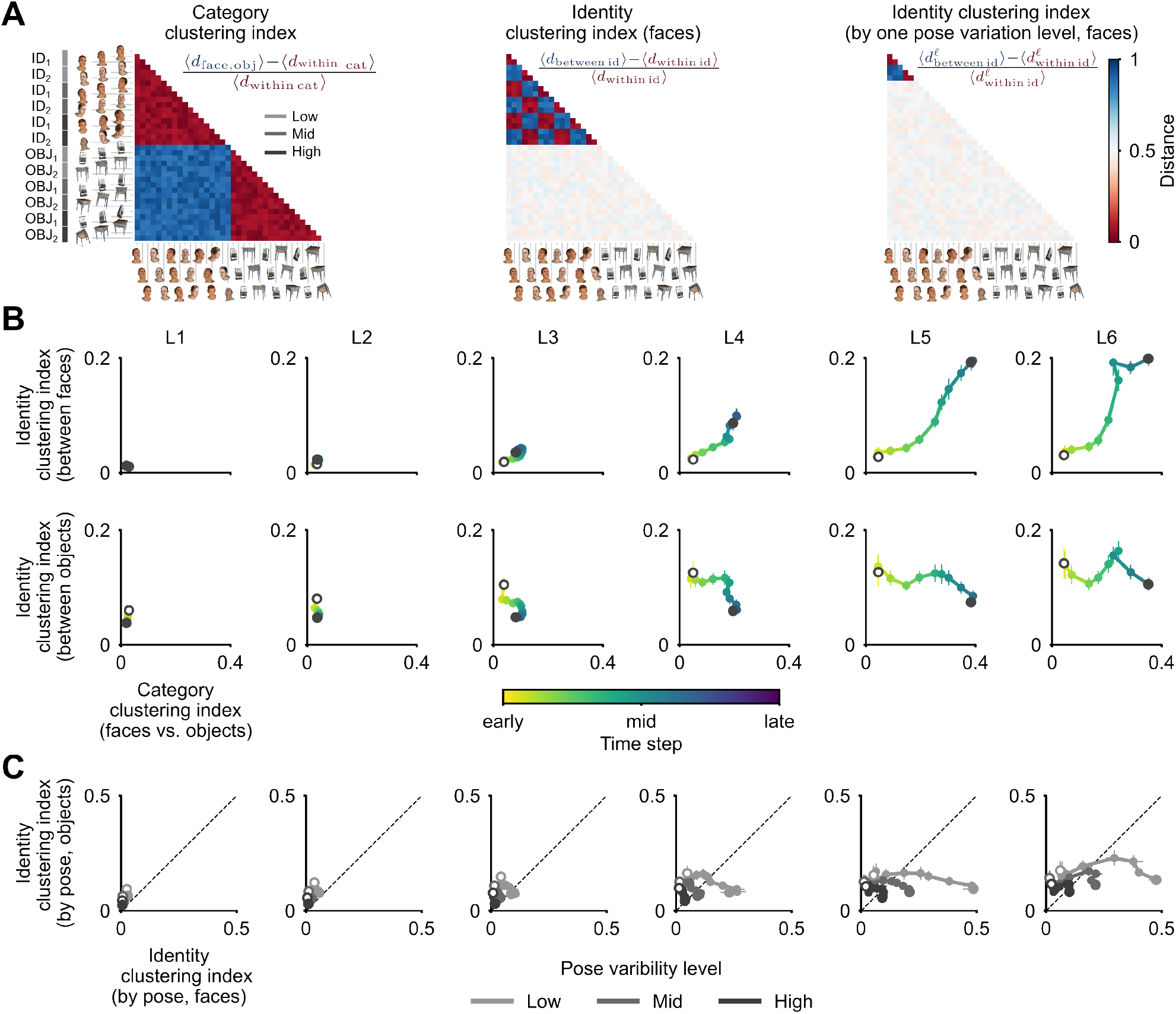
Geometry-derived category and identity discrimination metrics. All results shown are from the face-trained model. (**A**) Schematic illustration of the RDM-based metrics. Category discrimination is defined as the normalized difference between between-category (faces vs. objects) and within-category distances. Face-identity discrimination is defined as the normalized difference between between-identity and within-identity distances computed within the face category. Pose-conditioned identity discrimination further isolates pose-tolerant identity structure by comparing between-identity distances across pose variation with within-identity distances. Color indicates pairwise dissimilarity. (**B**) Joint evolution of category discrimination (*x*-axis) and identity discrimination (*y*-axis) across recurrent timesteps for each layer (L1–L6). The top row plots face-identity discrimination against category discrimination; the bottom row plots object-identity discrimination against category discrimination. Each point corresponds to one recurrent timestep and is colored from early to late. Across layers, category separation emerges by mid-layers and remains relatively stable, whereas identity discrimination strengthens primarily in deeper layers and at later recurrent timesteps. (**C**) Pose-conditioned identity discrimination for faces (*x*-axis) and objects (*y*-axis), shown separately for low, mid, and high pose-variation levels. Each trajectory corresponds to one variation level, with successive points tracing recurrent timesteps. In deeper layers, recurrence produces a substantially larger increase in pose-tolerant identity separation for faces than for objects, while preserving the relative ordering of the variation levels. Error bars indicate 95% bootstrap confidence intervals (1,000 resamples of identity-level block means).

Category discriminability was already established during the feedforward pass. Category clustering emerged in the middle layers (L3, L4) at intermediate timesteps and remained largely stable across recurrent timesteps. Identity clustering emerged in higher layers and at later timesteps (Fig. 5B). Lateral recurrence, thus, does not reorganize representations indiscriminately; it acts on the finer-grained distinctions the network was trained to resolve, increasing both selectivity to the task-relevant variables and tolerance to nuisance variation (in particular pose variation: yaw, pitch, and roll) (Rust and DiCarlo, 2010).

We further analyzed how recurrent processing affects identity clustering for different levels of pose variation (low, mid, high). For the face-trained model, we found that recurrent processing enhances identity clustering for all levels, but that clustering in the final representation is weaker the greater the variation of poses (Fig. 5C). This demonstrates that, while recurrence enhances identity clustering, it does not collapse different poses of the same face to a single point and remove pose information completely.

For the object-trained model, the overall pattern was complementary, with the trained domains reversed. Category structure again stabilized early, whereas recurrent processing selectively strengthened object-identity organization in deeper layers and at later timesteps, with little refinement of face-identity representations. This refinement persisted across pose-variation levels but remained weaker under greater pose variation. These population-level results leave open how selective refinement is implemented at the level of individual units. We therefore characterized the tuning dynamics of L6 units across recurrent timesteps (Fig. 6).

**Figure 6.**
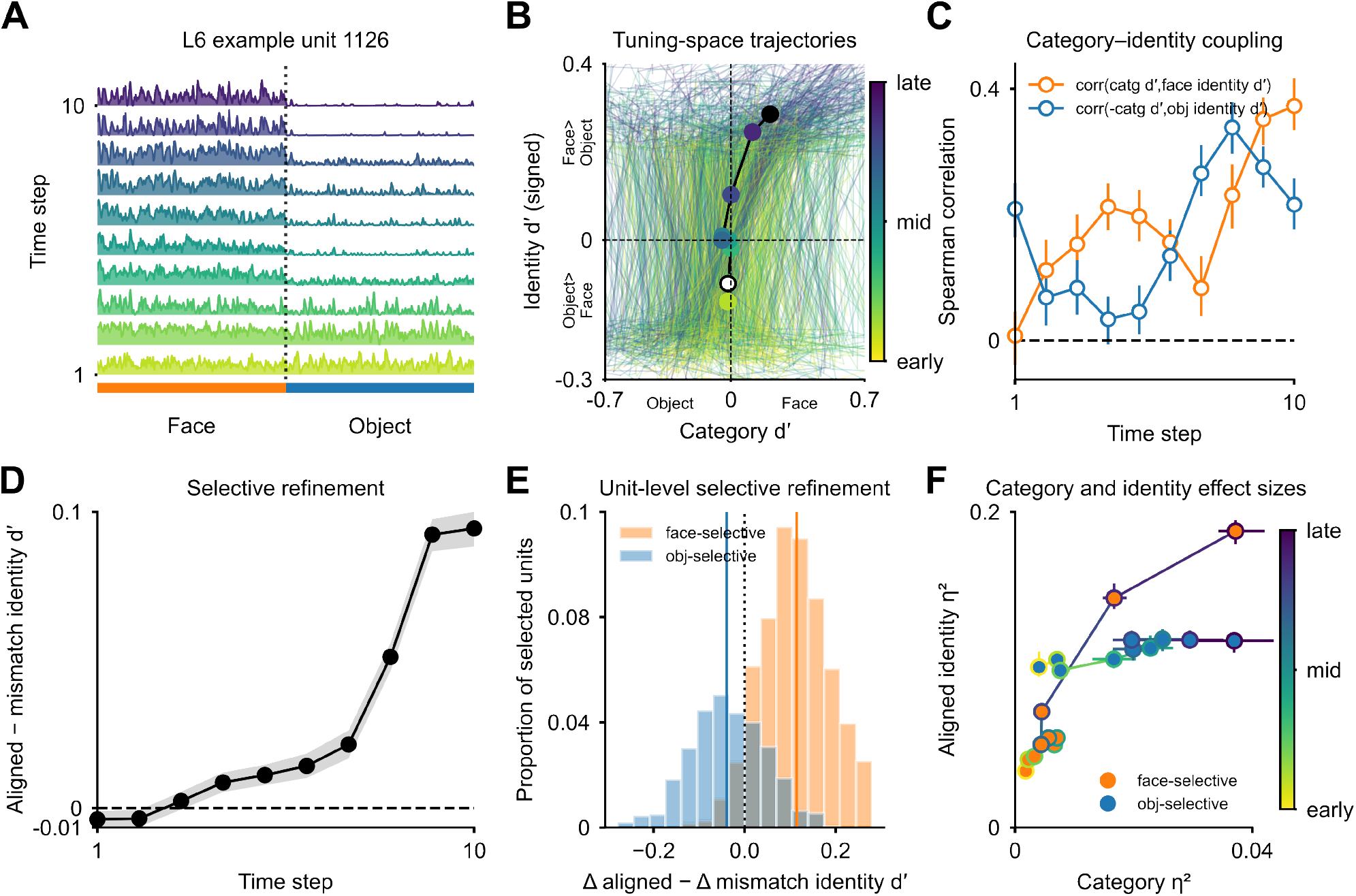
Recurrent computations strengthen both face-category selectivity and face-identity discrimination in the same units. All results shown are from the face-trained model. (**A**) Response profile of an example L6 unit (unit 1126) across recurrent timesteps in the face-trained model. Responses are shown for stimuli ordered by domain (faces, left; objects, right), with the dashed line marking the category boundary and traces colored by recurrent timesteps (early to late). (**B**) Trajectories of individual L6 units in a two-dimensional tuning space defined by category d′ (*x*-axis; negative, object-selective; positive, face-selective) and signed identity d′ (*y*-axis; positive, stronger face-identity discrimination; negative, stronger object-identity discrimination). Thin lines show trajectories for 500 randomly sampled units, and the bold line shows the average trajectory across all units. (**C**) Spearman correlations across L6 units between category d′ and face-identity d′ (orange), and between negative category d′ and object-identity d′ (blue), at each recurrent timestep. (**D**) Average selective-refinement index across all units over recurrent timesteps, defined as aligned minus mismatched identity d′, where aligned denotes the domain consistent with a unit’s category preference (each unit’s category preference was determined from category d′ computed on one half of the stimuli, while aligned and mismatched identity d′ values were measured on the held-out half). Shaded bands show bootstrap 95% confidence intervals. (**E**) Distribution of unit-level changes in selective refinement from early to late recurrence, computed as Δ aligned minus Δ mismatched identity d′, shown separately for face-selective (orange) and object-selective (blue) units. (**F**) Category and identity effect sizes, showing category *η*^2^ against aligned identity *η*^2^ for face-selective and object-selective units, with color indicating recurrent timesteps. Error bars show bootstrap 95% confidence intervals across units.

### Recurrence aligns category preference with identity sensitivity at the single-unit level

To understand how individual units contribute to the population-level refinement described above, we characterized the tuning dynamics of L6 units across recurrent timesteps in the face-trained model (Fig. 6). Individual units became more face-category selective across recurrent timesteps (Fig. 6A). We next represented each L6 unit’s state in a space defined by category d′ and signed identity d′, which we define as the larger of the two identity d′ values for faces and objects, with the sign positive if the face-identity d′ is larger and negative if the object-identity d′ is larger. Across recurrent timesteps (yellow to blue), units became more face-category-selective and more face-identity-discriminating. Each trajectory reflects how recurrent computation changes a single unit’s tuning. The increased density of the blue trajectories (reflecting high-stage L6 tuning) in the upper right demonstrates that the same units develop both face-category-selectivity and face-identity discrimination (Fig. 6B).

Across timesteps, the correlation between a unit’s category selectivity and its within-domain identity discriminability increased (Fig. 6C). We quantified this with a selective-refinement index: the difference between identity d′ in the category-aligned and category-mismatched domains, using cross-fitted splits (see Methods). In the face-trained model, this index increased steadily across recurrent timesteps (Fig. 6D). The distribution of unit-level changes confirmed that face-preferring units gained identity discrimination in their preferred domain (Fig. 6E). To assess how recurrence changes the variance associated with category and identity, we measured the effect size of each factor separately for every unit (Fig. 6F). Because *η*^2^ is a proportion of total variance, it is unchanged by any uniform rescaling of responses and registers only shifts in how variance is distributed across factors. At later timesteps, the fraction of response variance explained by identity increased in the category-aligned (face) domain, while the fraction explained by category was preserved. The added variance is specific in two respects: it loads on identity rather than category, and it appears in face-preferring units rather than across the population. This gives a unit-level mechanism for the population-level refinement.

The object-trained model showed the same signature with the domains reversed: over recurrent timesteps, object-selective units gained object-identity discrimination while preserving their category tuning, with weaker change in face-selective units (Supplementary Fig. S4). The mechanism is therefore not specific to faces. In both models, recurrence adds identity-related variance to the units already tuned to the trained category without diminishing their category selectivity.

## Discussion

### Lateral recurrence refines task-relevant representations

Our results show that lateral recurrence refines task-relevant representations over time within a neural population. For the same architecture and connectivity structure, varying the training objective (face versus object recognition) yielded a double dissociation: recurrence enhanced face-identity discrimination in the face-trained model, leaving object-identity discrimination unchanged, and vice versa in the object-trained model. This pattern suggests that lateral recurrence selectively refines task-relevant representations at higher levels. This does not preclude an additional domain-general role for recurrence at lower levels, although we did not see strong evidence of that here.

These results connect to longstanding debates about recurrent processing. Predictive coding and related frameworks have emphasized the domain-generality of recurrent processing, proposing that feedback and lateral interactions sharpen representations by suppressing predictable components largely regardless of content (Rao and Ballard, 1999; Lee and Mumford, 2003). Empirical studies using backward masking have similarly suggested that recurrence broadly strengthens visual representations across stimulus categories (Fahrenfort et al., 2007; Xie et al., 2025). Our results are about lateral connectivity only, but are broadly consistent with these observations: recurrence may well serve general denoising or stabilizing functions under degraded input conditions. We add evidence for a task-dependent component that operates alongside any generic mechanism, and that is visible only when the temporal evolution of representations is tracked across recurrent timesteps.

The perceptual learning literature showed that training enhances discriminability for practiced distinctions without globally sharpening untrained features (Schoups et al., 2001; Jehee et al., 2012). Perceptual learning likely depends on interactions between feedback signals and local cortical circuits, with effects determined by which intrinsic connections are engaged at any given time (Gilbert et al., 2009). Here, we identify a specific computational substrate for such selectivity: the temporal unfolding of lateral recurrence, which reshapes representational geometry within the same layer over successive iterations. Refinement does not require additional feedforward depth or architectural specialization; it emerges from the interaction between learned lateral weights and recurrent dynamics.

How identity refinement unfolds within a single face-selective area, rather than across the face patch hierarchy, has been difficult to study empirically. Face-selective regions in primate IT cortex carry both category and identity information, and identity representations become increasingly pose-tolerant across hierarchical stages (Freiwald and Tsao, 2010; Chang and Tsao, 2017), but the temporal trajectory within a single area has remained elusive. Our model explicates a computational hypothesis: that lateral recurrence supports within-area refinement shaped by the area’s training history. This offers a computational account of how domain-specialized regions sharpen their identity representations.

### A temporal division of labor: category emerges before identity

We observed rapid stabilization of category structure followed by expansion of the representational space occupied by different identities and a contraction of the variance due to nuisance variables. This dynamical computation is consistent with a classical result in primate neurophysiology. Sugase et al. (1999) showed that inferotemporal neurons first encode coarse category information and only later convey finer identity distinctions. Population-level decoding studies, similarly, showed that category information rapidly comes to be explicitly represented, with finer within-category distinctions becoming explicit later (Meyers et al., 2008; Dehaqani et al., 2016).

Explicit, here, means linearly decodable (Kriegeskorte and Douglas, 2019). By the data-processing inequality, new information cannot emerge from processing of the sensory input. All information present at a given stage must be present in each preceding stage of representation, including the input, but is not necessarily represented explicitly at lower levels. Because face identity differences are relatively subtle differences between physically similar objects (human faces) and because nuisance variables, such as pose and lighting, cause large differences in retinal appearance, neural manifolds corresponding to distinct faces are entangled in their low visual representations (DiCarlo and Cox, 2007). One way to render distinct identities linearly decodable is to reduce the representational variance associated with nuisance variables. In the macaque face-processing system, pose tolerance increases across hierarchical stages and time, and different poses of the same person’s face come to cluster in the representational space (Freiwald and Tsao, 2010; Chang and Tsao, 2017). Recurrent processing becomes especially important when feedforward computations are insufficient: in IT cortex, behaviorally sufficient solutions for images that are difficult to recognize emerge about 30 ms later than for control images (Kar et al., 2019). Time-resolved recordings across stages of ventral-stream processing show that representational geometry continues to evolve well beyond the initial feedforward sweep in humans and macaques (Kietzmann et al., 2019; Anthes et al., 2026). Our modeling results here suggest that lateral recurrence is among the mechanisms underlying these dynamic representational changes. Lateral recurrence expands between-identity distances relative to nuisance variation, while preserving the category partitions established during the feedforward pass.

The temporal division of labor, with categorical distinctions made explicit first and identity becoming explicit only later, changes how we should interpret recurrence-dependent improvements in recognition. If recurrence simply reduced noise and other low-level nuisance variation, then category and identity distances would grow simultaneously and by the same factor relative to that nuisance variation. Instead, the effect is selective: between-identity distances grow while the category partition is preserved, and this expansion is task-relevant: strongest in the trained domain. Recurrent iterations resolve within-category ambiguity that the feedforward sweep leaves unresolved. This is consistent with proposals that recurrence supports iterative repair of ambiguous or degraded representations (Tang et al., 2018), but adds an important constraint: what counts as “ambiguous” depends on the task the system was trained to solve.

In the face-trained model, what is initially ambiguous is the depicted person’s identity under pose variation. Different individuals remain distinctly represented (i.e., identity information is present), but pose spreads the images of a single identity such that they become entangled with images of other identities in the feed-forward representation and are not linearly separable (i.e., identity information is implicit). Recurrence reduces this nuisance variation and disentangles the identity representations, rendering the identity information explicit. In the object-trained model, what is initially ambiguous is object identity, and we see an analogous computational signature of dynamic disentangling, specific to the domain of objects.

Converging with this account, Shi et al. (2026) reported direct evidence for a closely related temporal division of labor in macaque face patches. Face-selective cells in inferotemporal cortex initially carried a general-purpose code, built from features shared with non-face objects and suited to face detection. Roughly 100 ms after onset, a rapid, concerted population event reconfigured this code into a face-specific one: response gradients along the detection-related dimensions reversed, and new tuning emerged for the finer features that distinguish individual faces. The dissociation is therefore temporal and category-specific: The system switches from an early code common to faces and other objects to a late code that discriminates among individual faces. This matches the prediction from our models: in domain-selective populations, recurrence widens the distances between identities of the preferred category and leaves between-identity distances for other categories approximately unchanged.

### A mechanistic account at the single-unit level

Our population-level analyses showed that recurrence clusters representations within each identity for the trained domain, but did not reveal how this clustering arises at the single-unit level. The single-unit results point to a mechanism. Recurrence does not increase identity sensitivity uniformly across units; the increase is concentrated in the units whose category preference matches the trained domain. In these units, category tuning is preserved while sensitivity to individual identities grows, so recurrence aligns each unit’s identity sensitivity with its category preference.

This alignment mechanism differs from two alternative accounts of how recurrent processing might improve recognition. Divisive normalization (Carandini and Heeger, 2012) is more naturally associated with a rescaling of responses than with a selective enhancement of one discriminative dimension over another. Attentional gain modulation (Reynolds and Heeger, 2009) could in principle produce task-relevant enhancement, but it requires an external attention field or top-down bias, neither of which exists in our architecture. The selectivity here is intrinsic to the learned lateral weights and emerges without external gating. The mechanism here is closer to accounts in which horizontal connections, feedback signals, and local microcircuits are shaped by experience and contribute to task-relevant processing (Roelfsema and Holtmaat, 2018; Gilbert and Li, 2013). Our results suggest a specific variant of this idea: the learned bias is a closer alignment between category preference and identity sensitivity, produced by the temporal dynamics of lateral interactions.

This account also fits recent circuit-level findings. In mouse primary visual cortex, Oldenburg et al. (2024) used targeted optogenetic stimulation of small neural ensembles to isolate the effect of recurrent activity from feedforward input. That effect depends jointly on cortical distance and feature preference: nearby co-tuned neurons are facilitated, while more distant or differently tuned neurons are suppressed. Znamenskiy et al. (2024) examined the synaptic basis of this specificity in the same area. Although inhibitory connectivity is dense, its strength is graded: one major class of inhibitory interneurons most strongly inhibits nearby excitatory cells with similar visual tuning. Recurrent interactions are therefore shaped by response similarity, not by proximity alone. Our single-unit results extend this principle to higher-order representations. In deeper layers of trained networks, lateral recurrence selectively increases identity sensitivity in units whose category preference aligns with the trained domain. Across the population, this widens the between-identity distances in the trained domain and improves cross-validated identity decoding there.

Our results have implications for the interpretation of mixed selectivity in ventral-stream populations. Mixed selectivity, where individual neurons carry information about multiple stimulus dimensions at once, is increasingly recognized as important for cortical computation (Rigotti et al., 2013; Fusi et al., 2016; Posani et al., 2026). Our results suggest that recurrence structures mixed selectivity by aligning category and identity dimensions according to the trained task. This structured mixing may be functionally useful. By concentrating identity sensitivity in units that already prefer the relevant category, the network co-locates category and identity information in the same units without the two trading off: category tuning is preserved as identity sensitivity is added, because category is carried by the coarse difference between preferred and non-preferred stimuli and identity by the finer variation within the preferred category. This should help downstream readout by category-selective populations.

### Limitations

The models were trained with backpropagation through time, which differs from known biological learning rules. However, the claims of this paper concern the functional consequences of learned lateral weights during inference, rather than the learning algorithm itself. Similar representational dynamics could emerge from biologically plausible learning rules that shape lateral connections to support the trained objective, including Hebbian or reward-modulated learning (Roelfsema and Holtmaat, 2018; Lillicrap et al., 2020). For the predictions developed here, what matters is the structure of the learned recurrent weights, not the learning algorithm.

The training pairings in our main networks couple content with task: face stimuli are paired with identification, object stimuli with categorization. Content and task therefore covary, and we cannot factorially separate their contributions in these networks. This pairing was deliberate. In natural perception, face recognition typically demands identifying the individual, and object recognition often merely requires a category label (Tanaka and Curran, 2001; Diamond and Carey, 1986). We chose to model the naturalistic alignment between domain and computational demand rather than treat content and task as independent factors. The jointly trained network (Supplementary Fig. S1), in which a single architecture is trained on both objectives simultaneously, provides partial dissociation: recurrence still refines identity selectively within each unit’s preferred domain, indicating that the selectivity does not require whole-network specialization for a single content/task pairing. Future work should include a more comprehensively controlled factorial design.

The two stimulus sets differ in internal homogeneity. Face identities were drawn from a single parametric shape and texture space and therefore differ only subtly, whereas the object exemplars come from distinct categories and differ along many dimensions at once; object-identity discrimination is closer to exemplar discrimination across heterogeneous types than to within-category individuation. This asymmetry cannot account for the main result, because the double dissociation reverses with the training objective while the stimuli are held fixed: the same object set is refined by the object-trained model and left largely unchanged by the face-trained model. It does mean the two domains are not matched in difficulty, so absolute decoding levels should not be compared across domains; the comparisons that carry the argument are within-domain, across timesteps and across models.

We focused on lateral recurrence because it can be isolated cleanly within a fixed architecture framework and dominates within-area connectivity, but the current study does not include inter-areal feedback or top-down attentional signals. Feedback connections likely implement additional computational mechanisms, for example injecting prior and contextual information, adjusting attentional modulation, or gating which domain benefits from recurrent refinement.

### Predictions and future directions

We use the models to derive predictions for time-resolved recordings in domain-specialized cortical areas.

The most direct prediction concerns the temporal dynamics of identity coding in face representations. Time-resolved recordings from face-selective patches in primate IT (Freiwald and Tsao, 2010; Chang and Tsao, 2017) should reveal that representational distances between identities expand later than those between categories, with this late expansion stronger for faces than for non-preferred object stimuli in the same region. Conversely, object-selective regions should show the opposite pattern. Existing multi-electrode datasets spanning the first 200–300 ms of processing may already contain the temporal signatures we predict, although the analyses would need to quantify category and identity geometry separately as a function of post-stimulus time. The roughly 30 ms delay in behaviorally sufficient solutions reported by Kar et al. (2019) is consistent with this prediction, though for object recognition rather than face identification. A direct test here would require within-region comparisons of category and identity decoding dynamics.

Perturbation experiments provide a complementary test. The graded relationship between recurrence strength and trained-domain decoding predicts that backward masking, cooling, or optogenetic perturbation of recurrent circuitry should impair pose-tolerant identity judgments more than rapid category discrimination (Fahrenfort et al., 2007; Kar and DiCarlo, 2021). This impairment should be domain-specific: disrupting recurrence in face-selective cortex should affect face-identity judgments more strongly than object-identity judgments presented in the same session.

The proposed mechanism also predicts a single-unit signature. During recognition, neurons with strong domain preference should become increasingly sensitive to identities within that domain while retaining their category preference. Implanted multi-electrode arrays could test this prediction by tracking the same neurons across early (*<*100 ms) and late (*>*150 ms) response epochs.

A convergent result across these tests would show that the temporal refinement seen in our models reflects a general property of domain-selective cortex, recurrence resolving within-category identity through the units already tuned to the relevant domain.

## Methods

### Stimulus generation

We generated two stimulus sets, faces and objects, each containing 24 identities rendered at three levels of pose variation (low, mid, high), with 12 images per identity at each level. This yielded 864 images per domain and 1,728 images in total.

#### Face stimuli

Face stimuli were generated using the FLAME parametric head model (Li et al., 2017) and an associated FLAME texture space (Karras et al., 2019; Feng and Bolkart). Each of the 24 identities was defined by a unique set of shape coefficients (300 dimensions) and texture coefficients (200 dimensions), sampled uniformly within ± 2 standard deviations of the corresponding model distributions. Expression parameters were fixed at zero to generate neutral faces. For each identity, 3D face geometry was obtained by passing shape parameters through FLAME, yielding a mesh with UV texture coordinates. Texture maps were reconstructed by combining the mean texture with identity-specific coefficients over the texture basis principal components. FLAME’s internal jaw/neck pose parameters were fixed at neutral; the yaw/pitch/roll variation described below was applied at the rendering stage.

#### Object stimuli

Object stimuli consisted of 24 3D object models obtained from Poly Haven, a library of high-quality 3D assets released under a CC0 license (https://polyhaven.com). Objects were selected to maximize category diversity and visual distinctness. Accordingly, object-identity decoding should be interpreted as exemplar discrimination across heterogeneous object types within the broader object class.

#### Pose variation

We defined three levels of pose variation by the ranges of yaw, pitch, and roll applied to each identity: low (±30° yaw, ±15° pitch, ±5° roll), mid (±60° yaw, ±30° pitch, ±10° roll), and high (±90° yaw, ±45° pitch, ±20° roll). For each identity and variation level, 12 poses were sampled within the specified ranges.

#### Rendering pipeline

All stimuli were rendered at 512 × 512 pixels in `pyrender` using a perspective camera with a 40° field of view and a 1:1 aspect ratio. Illumination was provided by a standard three-point lighting setup, together with low ambient light.

Meshes were centered at the bounding-box centroid and scaled so that their largest dimension matched a target size of 1 scene unit. Camera distance was then adjusted to control image framing, with tighter framing for faces and wider framing for objects.

Rendered RGBA images were composited onto natural-image backgrounds that were randomly sampled from 1,728 images across six scene categories from the Intel Image Classification dataset (Intel and Puneet6060, 2018).

### Model architecture and training

We used the BL (bottom-up + lateral) variant of the recurrent BLT architecture (Spoerer et al., 2017, 2020). The network comprised four convolutional (L1–L4) and two fully connected layers (L5–L6) with two connection types: bottom-up (feedforward) connections between successive layers and lateral (within-layer) recurrent connections. Lateral connections were implemented as within-layer convolutional kernels, conv_*i→i*_, allowing recurrent dynamics to unfold over discrete timesteps while preserving the spatial resolution of the feedforward pathway. Top-down (feedback) connections, present in the full BLT architecture, were not included; the models studied here thus isolate the contribution of lateral recurrence specifically.

We compared two models with identical architectures but different training objectives. The *face-trained model* was optimized for face identification using the VGGFace2 dataset (Cao et al., 2018) (3,890 identities), and the *object-trained model* was optimized for object categorization using ImageNet (Deng et al., 2009) (1,000 categories). Training images were matched at 1.28 million per model. For VGGFace2, we used the first 3,890 identities in directory order (selected deterministically for reproducibility); for ImageNet, we used all 1,000 categories. Both models were trained with backpropagation through time for 15 recurrent timesteps, with a linear readout head on the final layer, using Adam (Kingma and Ba, 2014) with batch size 64, learning rate 5 × 10^−4^, and weight decay 10^−4^ for 100 epochs. The object-trained model achieved 55% top-1 accuracy on ImageNet and the face-trained model achieved 83% top-1 accuracy on VGGFace2.

To partially dissociate content from task, we trained a third network with the same architecture on both objectives simultaneously. Two linear readout heads were attached to L6, one for face identification (VGGFace2, 3,890 identities) and one for object categorization (ImageNet, 1,000 categories).

### Biological unrolling

Following Kubilius et al. (2018) and Spoerer et al. (2020), we adopted a staggered unrolling scheme in which bottom-up and recurrent connections each take one discrete timestep. The computational graph therefore reflects the propagation delay of the feedforward sweep: each layer receives its first input only after the feedforward signal has arrived from the previous layer. As a result, the number of available recurrent timesteps differed across layers: layer 1 received input at all 15 timesteps, while layer 6 received input starting from timestep 6, yielding 10 available timesteps for analysis (Fig. 3).

### Image preprocessing and feature extraction

All stimulus images were resized to 224 × 224 pixels and normalized with ImageNet channel statistics before being passed to the models. For each recurrent model, a forward pass yielded activations at all six layers across recurrent timesteps. Spatial activation maps were flattened to produce one feature vector per image and timestep, yielding a feature matrix **X** ∈ R^*N×D*^ for each layer–timestep combination, where *N* = 1,728 is the total number of stimuli and *D* is the number of units.

### Representational dissimilarity analysis

We computed representational dissimilarity matrices (RDMs) for each layer–timestep combination and compared them using representational similarity analysis (Kriegeskorte et al., 2008). RDMs were computed for each layer–timestep combination from the corresponding feature matrices. Pairwise distances were computed as 1 −Pearson correlation (or 1 −Spearman correlation for the second-order analysis; Euclidean distance for the supplementary control, Supplementary Fig. S3). The resulting *N × N* symmetric distance matrices were used for all geometry-based analyses.

#### RDM-derived discrimination indices

To quantify the dis-criminative structure embedded in the representational geometry, we derived three scalar indices from each RDM.

*Category clustering index* was defined as the normalized difference between the mean between-category distance and the mean within-category distance:

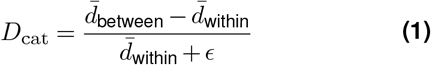

where *ϵ* is a small constant added for numerical stability.

*Identity clustering index* (computed separately for faces and objects) was defined as the normalized difference between mean between-identity and mean within-identity distances, computed within a single category:

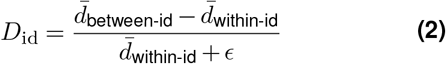

*Pose-conditioned identity clustering index* the identity index was recomputed separately within each pose-variation level, using identity blocks restricted to that level.

Standard errors for all indices were estimated using bootstrap resampling (1,000 iterations) over identity-level block means, resampling identities with replacement while preserving the within-identity sample structure.

#### Second-order representational similarity analysis

To quantify how representational geometry changes across layers and timesteps, we computed second-order distances between all pairs of RDMs. Each RDM was vectorized (upper triangle), and pairwise Spearman rank correlation distances (1™*ρ*) were computed between all layer–timestep combinations, yielding a second-order distance matrix. This matrix was visualized using classical multidimensional scaling (MDS) to embed each layer–timestep state as a point in two dimensions, revealing the trajectory of representational change across the model hierarchy and recurrent time.

### Cross-validated linear decoding

Linear decodability was evaluated with logistic-regression classifiers applied to the feature vectors at each layer and timestep. For each cross-validation fold, features were standardized using parameters (mean and variance) computed on the training split and then applied to the held-out data. We performed 24-way classification of face (or object) identity using a leave-one-level-out cross-validation scheme across pose variation levels. In each fold, classifiers were trained on images from two pose levels and tested on the remaining level, ensuring that performance reflects pose-tolerant identity representations. Decoding accuracy was computed for each fold and then averaged across the three levels. Decoders used the L-BFGS solver with a maximum of 1,000 iterations.

### Parametric gating of lateral recurrence

To test whether identity decoding causally depends on lateral recurrence, we parametrically manipulated recurrence strength by scaling the weights of all lateral convolutional kernels (conv_*i*_→_*i*_ for *i* = 1,…, 6) by a scalar gate value *g* ∈ {1, 0.75, 0.5, 0.25, 0}, where *g* = 1 corresponds to the intact model and *g* = 0 corresponds to full ablation of lateral connections. For each gate value, all lateral weights were multiplied by *g*, and the full analysis pipeline (feature extraction, identity decoding) was rerun. Decoding results were evaluated at three temporal durations (*T* = 2, *T* = 6, *T* = 10) to assess whether the dependence on recurrence strength scales with temporal depth.

### Single-unit selectivity metrics

All single-unit analyses were performed on units in the final layer (L6). For each unit, responses were defined as the scalar activation value for each stimulus, and the following metrics were computed at each recurrent timestep:

*Category d*′. A signed sensitivity index quantifying the degree to which a unit discriminates faces from objects:

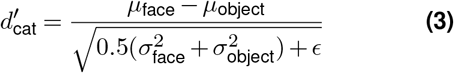

where *µ* and *σ*^2^ denote the mean and variance of unit

Farzmahdi *et al*. | Task-dependent recurrent refinement responses across all stimuli in each category. Positive values indicate face selectivity; negative values indicate object selectivity.

*Identity d*′. For each unit, identity *d*′ was computed for all pairwise identity comparisons within a given category (face or object) using the same formula applied to the distributions of responses for each identity. The mean over all pairwise comparisons yielded an average identity *d*′ per unit, separately for faces and objects. The larger of the two values was retained, with the sign indicating whether face-identity *d*′ exceeded object-identity *d*′ (positive) or vice versa (negative).

#### Selective-refinement index

For each unit, the selective-refinement index was defined as the difference between identity *d*′ in the category-aligned domain (matching the unit’s category preference) and identity *d*′ in the category-mismatched domain. In this analysis, category preference and identity *d*′ were measured on independent splits of the stimulus set: stimuli were partitioned into two halves balanced by category, identity, and pose-variation level; category *d*′ was computed on the first split to determine each unit’s category preference (positive: face-preferring; negative: object-preferring); aligned and mismatched identity *d*′ values were then computed on the held-out second split. Results were averaged across both partition assignments. An increase in the selective-refinement index over recurrent time indicates that the unit is enhancing identity coding for the domain matching its category preference, computed independently of the data used to define that preference.

#### Category–identity alignment

Category–identity alignment was quantified across all L6 units at each recurrent timestep. For faces, we computed the Spearman correlation between category *d*′ and face-identity *d*′. For objects, we computed the correlation between negative category *d*′ and object-identity *d*′. Thus, a positive correlation indicates greater identity sensitivity in units preferring the corresponding domain.

### Factor-specific effect-size analysis

To quantify how strongly category and identity were associated with unit responses across recurrent timesteps, we computed *η*^2^ separately for each grouping factor using a one-way ANOVA. For each L6 unit at each recurrent timestep, we computed:

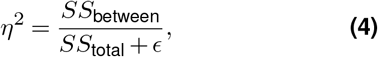

where *SS*_between_ is the sum of squares between the groups defined by the factor and *SS*_total_ is the total sum of squares. Three grouping factors were analyzed separately: category (face vs. object, computed across all stimuli), face identity (24-way, computed across face stimuli), and object identity (24-way, computed across object stimuli). To assess category–identity alignment, each unit’s category *η*^2^ was plotted against its aligned identity *η*^2^—face-identity *η*^2^ for face-selective units and object-identity *η*^2^ for object-selective units, at each recurrent timestep.

### Statistical procedures

Decoding accuracy is reported as the mean across the three leave-one-level-out cross-validation folds (one fold per pose-variation level), with error bars showing the standard error across folds (*n* = 3). Recurrence gain, defined as the difference between decoding accuracy at the final and initial recurrent timesteps within each pose-variation level, is reported with the analogous fold-wise SEM.

For RDM-derived discrimination indices, error bars indicate bootstrap standard errors over identity-level block means (1,000 resamples).

For single-unit analyses, error bars indicate 95% bootstrap percentile confidence intervals obtained by resampling units with replacement (1,000 resamples).

All resampling used a fixed seed. We did not apply a multiple comparisons correction: estimates across layers, timesteps, and conditions are dependent and address a single hypothesis rather than separate confirmatory questions. Comparisons between conditions (trained vs. non-trained domain; first vs. last timestep; intact vs. gated) are reported as effect sizes with the uncertainty estimates defined above. Our conclusions rest on the magnitude of these effects and on the consistency of the pattern across decoding, representational geometry, recurrence-gating, and single-unit analyses, rather than on any individual comparison.

## ACKNOWLEDGMENTS

Research reported in this publication was supported by the National Institute of Neurological Disorders and Stroke of the National Institutes of Health under Award Number RF1NS128897.

## DATA AVAILABILITY

The dataset will be made publicly available upon publication at doi:10.5281/zenodo.22101312.

## Supplementary Information

**Figure S1.**
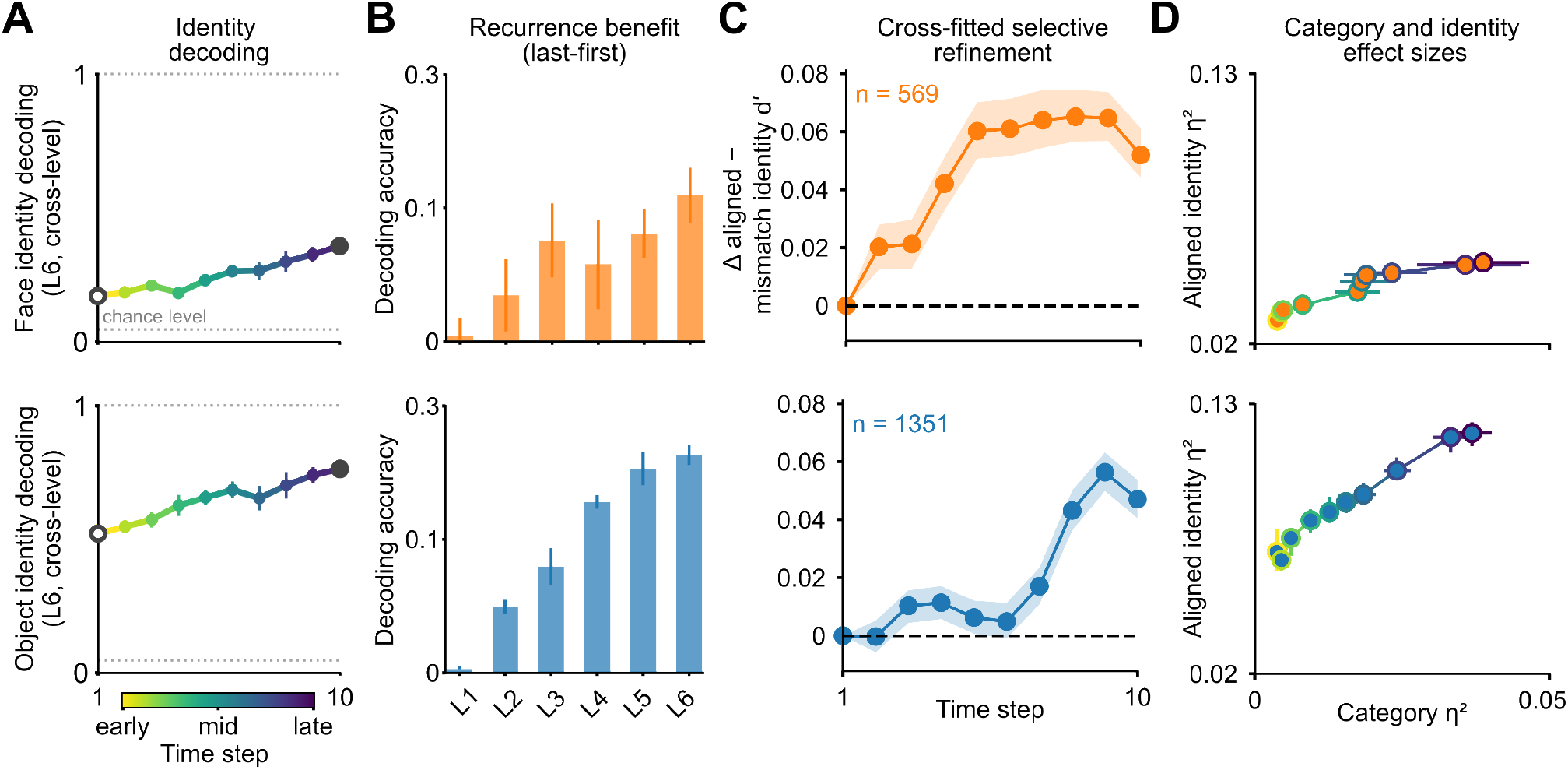
Population- and unit-level analyses of recurrent processing in the jointly-trained model. Analyses for the model trained on both faces (VGGFace2) and objects (ImageNet). Top row shows results for face-selective units; bottom row shows results for object-selective units. (**A**) Population decoding across recurrent time. Curves show decoding performance for the corresponding preferred domain across recurrent steps; points are colored by time step (early to late, color bar below). The dotted horizontal line indicates chance level. (**B**) Population recurrent gain across layers, computed as the late-minus-early change in decoding accuracy for each layer. Bars show mean gain and error bars indicate SEM across leave-one-level-out viewpoint splits. (**C**) Cross-fitted single-unit selective-refinement time course. Units were first classified as face- or object-selective using category *d*′ computed on one split of the data, and aligned and mismatched identity *d*′ were then measured on the held-out split. Selective refinement was defined as aligned minus mismatched identity *d*′, where aligned identity *d*′ corresponds to face identity *d*′ for face-selective units and object identity *d*′ for object-selective units. Values are plotted as the change relative to the first recurrent time point, so positive values indicate an increase over time in identity sensitivity aligned with the unit’s category preference. Lines show the mean across units, shaded bands show bootstrap 95% confidence intervals, and the number of stable units included in each analysis is indicated in each panel. The smaller number of face-selective units reflects the higher class count and harder optimization landscape of fine-grained face identification relative to object categorization. (**D**) Single-unit variance decomposition for the same stable units. Trajectories plot category *η*^2^ against aligned identity *η*^2^ across recurrent time, with points colored by time step. Error bars show bootstrap 95% confidence intervals across units. Together, these analyses show that recurrent processing strengthens category-aligned identity information in both face-selective and object-selective subpopulations within a single jointly-trained network. The same coupling between category preference and identity refinement therefore emerges within one model and one training regime, generalizing the dissociation observed in the single-task models.

**Figure S2.**
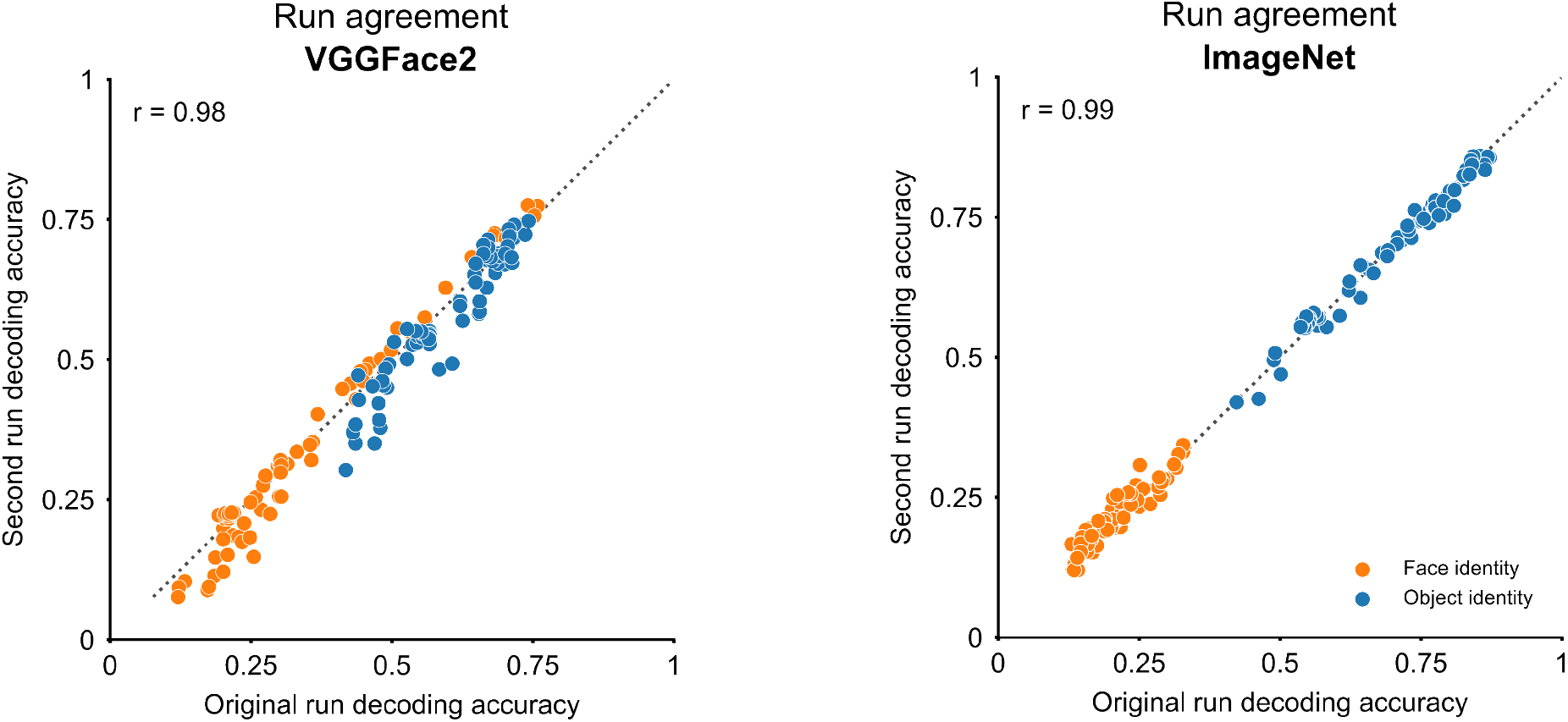
Reproducibility of decoding results across independent training runs. Decoding accuracy from the original training run (x-axis) is plotted against decoding accuracy from a second independent training run with a different random initialization (y-axis), separately for the VGGFace2-trained model (left; r=0.98) and the ImageNet-trained model (right; r=0.99). Each point corresponds to one (layer × recurrent timestep) combination (n=75). Points fall close to the unity line (dotted), indicating that the patterns reported in the main text are stable across independent training runs. Orange: face-identity decoding; blue: object-identity decoding.

**Figure S3.**
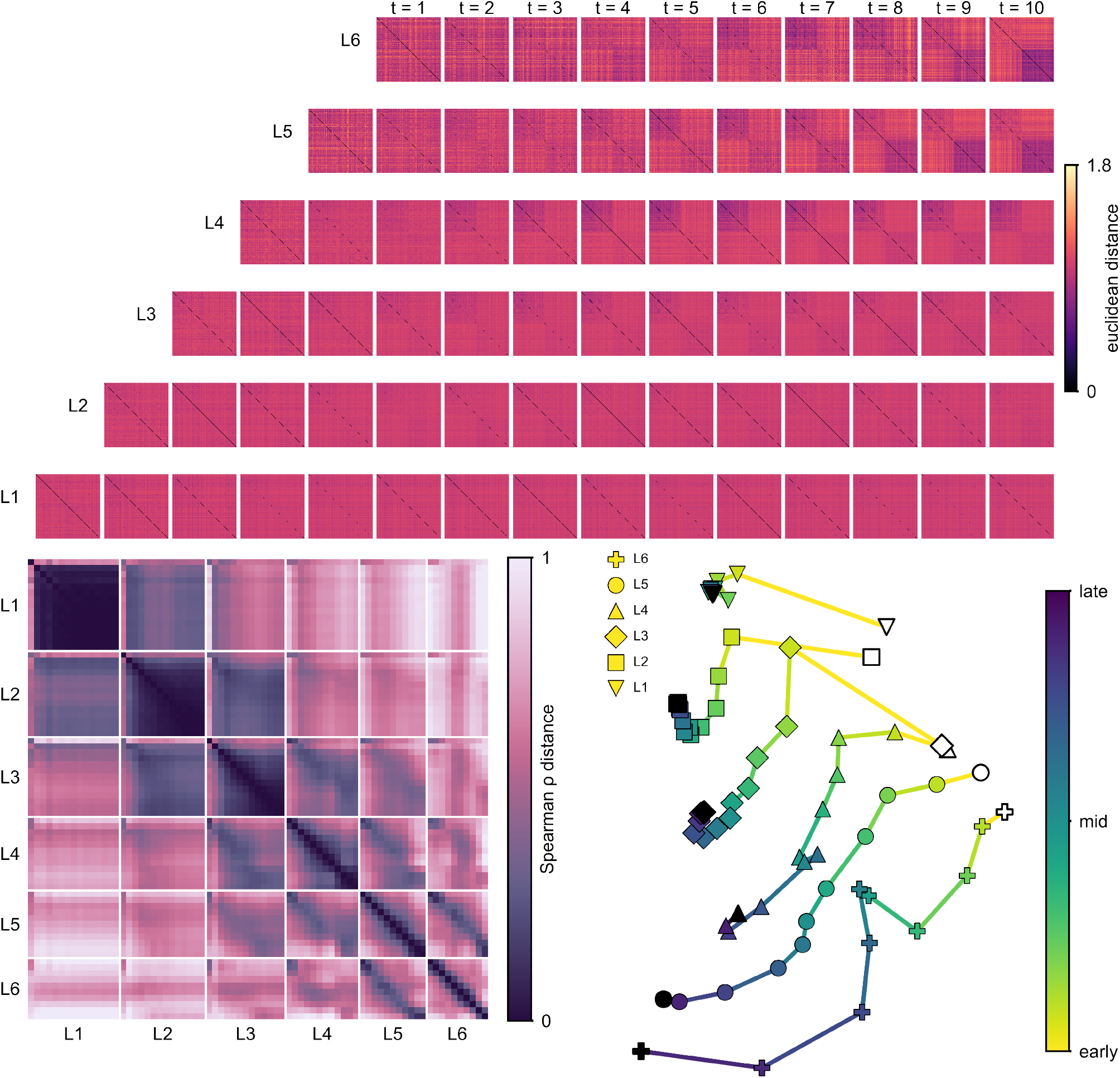
Population geometry results replicate under Euclidean distance. Identical analyses to Figure 3, but with first-order representational dissimilarity computed using normalized Euclidean distance rather than correlation distance. Pairwise Euclidean distances were computed between population response vectors, normalized by the square root of the number of units in the layer to control for differences in dimensionality, and then divided by the mean off-diagonal distance within each RDM so that color values represent dissimilarity relative to each RDM’s average. Color values therefore indicate within-RDM relative structure rather than absolute distances, which would otherwise be confounded by timestep-to-timestep fluctuations in overall response magnitude inherent to recurrent dynamics. (**A**) RDMs across layers and recurrent timesteps, organized as in Figure 3A. (**B**) Second-order Spearman *ρ* distance matrix between RDMs from each layer–timestep combination. (**C**) MDS embedding of the second-order distance matrix. The temporal evolution of representational geometry, the stability of early-layer representations, and the larger displacement of mid-to-late layers across recurrent time all replicate the patterns observed under correlation distance (Fig. 3), confirming that the main conclusions are robust to the choice of distance metric.

**Figure S4.**
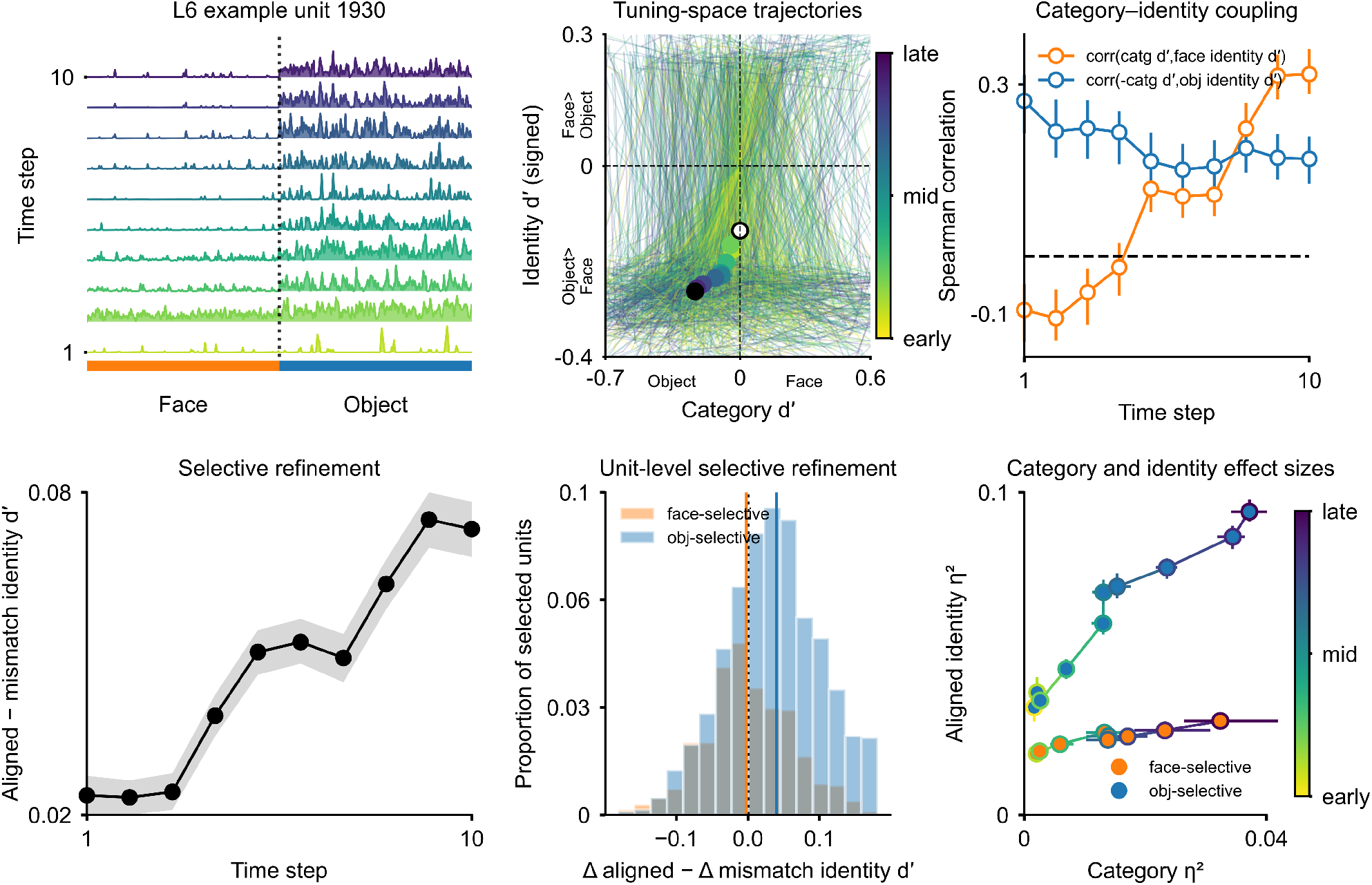
Recurrent computations strengthen both object-category selectivity and object-identity discrimination in the same units in the object-recognition-trained model. Panels follow the same layout as Fig. 6, computed here for the object-trained model. (**A**) Response profile of an example L6 unit (unit 1930) across recurrent timesteps. (**B**) Trajectories of individual L6 units in the tuning space defined by category d′ and signed identity d′. (**C**) Spearman correlations across L6 units between category d′ and face-identity d′ (orange), and between negative category d′ and object-identity d′ (blue), at each recurrent timestep. (**D**) Average selective-refinement index across units (aligned minus mismatched identity d′; cross-fitted, see Methods). (**E**) Distribution of unit-level changes in selective refinement from early to late recurrence, for face-selective (orange) and object-selective (blue) units. (**F**) Variance decomposition, plotting category *η*^2^ against aligned identity *η*^2^, colored by recurrent timestep. Across timesteps, object-selective units gain object-identity discrimination while preserving category tuning, mirroring the face-trained model with the domains reversed. Conventions and statistics as in Fig. 6.

